# *Pseudomonas putida* chromosomal toxin-antitoxin systems carry neither clear fitness benefits nor big costs

**DOI:** 10.1101/2020.03.18.996504

**Authors:** Sirli Rosendahl, Hedvig Tamman, Age Brauer, Maido Remm, Rita Hõrak

## Abstract

Chromosomal toxin-antitoxin (TA) systems are widespread genetic elements among bacteria, yet, despite extensive studies in the last decade, their biological importance remains ambivalent. The ability of TA-encoded toxins to affect stress tolerance supports the hypothesis of TA systems being associated with stress adaptation. However, the deletion of TA genes has usually no fitness consequences, supporting the selfish elements hypothesis. Here, we aimed to evaluate the cost and benefits of chromosomal TA systems to *Pseudomonas putida*. We show that multiple TA systems do not confer fitness benefits to this bacterium as deletion of 13 TA loci does not influence stress tolerance, persistence and biofilm formation. Our results instead show that TA loci are costly and decrease the competitive fitness of *P. putida*. Still, the cost of multiple TA systems is low and detectable in certain conditions only. Construction of antitoxin deletion strains showed that only five TA systems code for toxic proteins, while other TA loci have evolved towards reduced toxicity and encode non-toxic or moderately potent proteins. Analysis of *P. putida* TA systems’ homologs among fully sequenced *Pseudomonads* suggests that the TA loci have been subjected to purifying selection and that TA systems spread among bacteria by horizontal gene transfer.

## Introduction

Bacterial chromosomes contain multiple copies of toxin-antitoxin (TA) gene pairs which code for a toxic protein and an antagonist of the toxin. Chromosomal TA systems most probably originate from plasmids^1-3^, where they contribute to plasmid maintenance during host replication^4^. The mechanism behind the plasmid stabilization by TA modules is the unequal stability of the toxin and antitoxin, with the latter being usually less stable than the toxin^5,6^. Loss of plasmid and termination of antitoxin production therefore results in release of toxin from the antitoxin-mediated control which ultimately leads to toxin-caused cell death or growth arrest and plasmid-free bacteria are outcompeted from the population^7,8^.

While the role of TA systems in plasmid addiction is highly recognized, the biological importance of chromosomal TA loci has remained enigmatic despite years of extensive research. Several functions have been attributed to the chromosomal TA systems including stabilization of genomic mobile elements^9,10^, defence against bacteriophages^11,12^ as well as other invading mobile DNA^13^ and modulation of bacterial stress tolerance^14^. However, there are also a lot of contradictions and discrepancy between different studies. For example, chromosomal TA loci were considered to be the major players in formation of metabolically dormant and antibiotic tolerant persister cells but recently this concept was rebutted^15-17^. Also, despite multiple studies show that TA toxins can affect the stress tolerance^18-22^, these studies mostly involve artificial overexpression of the toxin which will likely never occur from the single-copy chromosomal TA locus. It is also important to note that even though several studies report that chromosomal TA loci are upregulated by stress conditions^17,23,24^, it does not necessarily mean that toxin is liberated from the antitoxin-mediated control, particularly, when considering that antitoxin is produced at a much higher rate than the toxin^25,26^.

The potential benefit of chromosomal TA loci is even more ambiguous, given that deletion of the whole TA locus has usually no effect on bacterial fitness^20,27-31^. It has been therefore debated that multiple TA systems may contribute to bacterial stress tolerance synergistically. The evidence in favour of this hypothesis can be found in *Mycobacterium tuberculosis* where deletion of three chromosomal *mazF* genes impaired the survival of Mtb under oxidative stress, in nutrient-limiting conditions and in macrophages^32^. However, studies in *Escherichia coli, Salmonella enterica* and *Staphylococcus aureus* show that deletion of 10, 12 or 3 chromosomal TA loci, respectively, do not influence either tolerance to antibiotics or persister formation^15,16,33,34^.

*Pseudomonas putida*, a model soil bacterium known for its metabolic versatility and remarkable environmental adaptability^35^, is predicted to contain up to 15 chromosomal TA operons^36^. Four of these TA loci – *graTA, mqsRA, res-xre* and *mazEF* – are verified to comprise functional TA proteins. The most thoroughly studied is HigBA family GraTA module, which codes for an unusually stable antitoxin GraA^37^ and a ribosome-dependent mRNase GraT characterized by conditional toxicity^28,38^. Deletion of antitoxin gene *graA* from *P. putida* chromosome affects bacterial growth at 30 °C only slightly, but causes severe growth and ribosome biogenesis defect at lower temperatures^28,38-40^. GraT has ability to affect stress tolerance both positively and negatively, yet deletion of the *graTA* operon has no effect on *P. putida* fitness^28^. *P. putida mqsRA* locus also codes for a quite mild toxin, given that the removal of the antitoxin gene *mqsA* does not significantly affect cell growth^41^. Still, the MqsR toxin has retained some growth suppression activity, as deletion of the antitoxin resulted in increased formation of ciprofloxacin tolerant persister cells^41^. However, akin to *graTA*, the absence of the whole *mqsRA* locus did not affect the persister phenotype^41^. RES toxin of the RES-Xre module is structurally related to NADase toxins and its overexpression results in growth suppression due to NAD^+^ depletion^20,42^. Unfortunately, the consequences of the chromosomal deletion of *res-xre* genes to *P. putida* were not examined^42^. The *P. putida* MazEF system has been also analysed only in mechanistic sense in a study demonstrating that the purified MazF acts as an mRNase which cleaves UAC sequence^43^.

Given that TA systems code for toxic proteins it is reasonable to hypothesize that TA loci incur fitness costs to the bacterium whenever they enter to the chromosome via horizontal gene transfer. While great amount of studies on chromosomal TA systems deal with deciphering their putative biological roles, less attention is paid to the fitness costs the bacteria may pay for possessing multiple TA loci. Here we aimed to analyse the fitness benefits and costs of multitude TA systems to *P. putida*. To this end, 13 TA loci were deleted from the genome of *P. putida* strain PaW85 and stress tolerance, biofilm and persister formation as well as competitive fitness of Δ13TA strain was compared with that of wild-type. Our data indicate that TA systems do not provide a detectable advantage to *P. putida*. TA loci rather impose a burden to the bacterium because freeing *P. putida* of TA loci slightly increased the competitive fitness of the deletion strain. Deletion analysis of antitoxins revealed that five out of 13 TA loci encode toxic proteins, while others code for non-toxic or only mildly potent proteins. Bioinformatic analysis revealed highly sporadic distribution of *P. putida* TA loci homologs in other *Pseudomonads* indicating that TA modules have been acquired horizontally.

## Results

### Construction of *P. putida* devoid of 13 TA systems

According to the TADB2 database, *P. putida* KT2440 is predicted to have 15 chromosomal TA operons^36^. In order to analyze the benefits and costs of multitude TA systems, the 13 TA loci were deleted from *P. putida* PaW85 (isogenic to KT2440) chromosome in the following order: *graTA* (PP_1586-1585), *res-xre* (PP_2433-2434), *higBA* (PP_1199-1198), *hicAB-1* (PP_1480-1479), *relE*_*2*_*-higA*_*2*_ (PP_5435-PP_0274), *mqsRA* (PP_4205-4204), *mazEF* (PP_0770-0771), PP_1716-1717, *brnTA* (PP_4530-4529), PP_4151-4152, *relBE* (PP_1268-1267), *yefM-yoeB* (PP_2940-2939) and *relB*_*2*_*-parE* (PP_2499-2500) (Table 1). The functionality of only four (*graTA, res-xre, mqsRA* and *mazEF*) out of these 13 TA modules has been studied experimentally^28,38,39,41-43^, while the other loci are putative TA pairs predicted *in silico* only^36^. TADB2 database annotates two more putative TA loci, the *hicAB-2* (PP_3900-3899) and PP_3032-3033, which are located in prophage 1 and prophage 2 genomes, respectively^36,44^. We kept these prophage-located TA loci in *P. putida* Δ13TA strain intact because attempting to delete the *hicAB-2* resulted in loss of prophage 1 (unpublished data). As the prophage loss is known to affect the fitness of *P. putida*^44^, this could complicate the interpretation of phenotypes resulting from deletion of other TA systems.

**Table 1.** 13 TA systems of *P. putida*

| Nr. | Locus tag* | Gene names* | Toxin's predicted function | Antitoxin deletion strain | Toxin toxicity | Homologs in pseudomonads | Toxin gene missing or truncated |
| --- | --- | --- | --- | --- | --- | --- | --- |
| 1 | <u>PP_1586-1585</u> | <u>graT-graA</u> | Inhibits ribosome biogenesis, ribosome-dependent mRNase | ΔgraA | | 180 | 16 |
| 2 | <u>PP_2433-2434</u> | xre-<u>res</u> | Reduces intracellular NAD<sup>+</sup> levels | Δxre | | 46 | 24 |
| 3 | <u>PP_1199-1198</u> | <u>higB-higA</u> | Ribosome-dependent mRNase | - |  | 37 | 10 |
| 4 | <u>PP_1480-1479</u> | <u>hicA1-hicB1</u> | Ribosome-independent mRNase | ΔhicB | | 41 | 6 |
| 5 | <u>PP_5435-0274</u> | <u>relE<sub>2</sub>-higA<sub>2</sub></u> | Ribosome-dependent mRNase | - |  | 8 | 3 |
| 6 | <u>PP_4205-4204</u> | <u>mqsR-mqsA</u> | Ribosome-independent mRNase | ΔmqsA | | 45 | 7 |
| 7 | <u>PP_0770-0771</u> | <u>mazE-mazF</u> | Ribosome-independent mRNase | ΔmazE | | 33 | 2 |
| 8 | <u>PP_1716-1717</u> | | | Δ1716 | | 332 | 2 |
| 9 | <u>PP_4530-4529</u> | <u>brnT-brnA</u> | Ribosome-independent mRNase | ΔbrnA | | 43 | 10 |
| 10 | <u>PP_4151-4152</u> |  |  | - |  | 72 | 6 |
| 11 | <u>PP_1268-1267</u> | relB-<u>relE</u> | Ribosome-dependent mRNase | ΔrelB | | 6 | 0 |
| 12 | <u>PP_2940-2939</u> | yefM-<u>yoeB</u> | Ribosome-dependent mRNase | - |  | 44 | 1 |
| 13 | <u>PP_2499-2500</u> | relB<sub>2</sub>-<u>parE</u> | Ribosome-dependent mRNase | - |  | 18 | 1 |

\*- Toxin underlined; ■ - Toxic, deleting the toxin gene is impossible; ■ - Moderately toxic, affects growth rate and stress tolerance; ■ - Not toxic, but affects stress tolerance in some conditions; ■ - Not toxic in any tested conditions

Whole-genome sequencing of *P. putida* wild-type strain PaW85 and Δ13TA showed that sequential deletion of TA loci has not caused accumulation of mutations in other regions of genome. Besides TA deletions we only detected 23 point mutations in Δ13TA that were not found in wild-type *P. putida* PaW85 (Table S1). 19 of these differences were located in intergenic regions, pseudogenes or in transposase genes. Comparison of *P. putida* PaW85 with KT2440 revealed 49 point mutations between the two strains, which, again, are mostly located in intergenic regions (Table S1).

To analyze the consequences of the lack of 13 TA systems at the whole-cell level, the proteomes of wild-type and Δ13TA strains grown in LB medium to mid-exponential phase were compared. Analysis of 1867 proteins detected in all three replicates of both strains showed that none of them were significantly differentially expressed between strains (Fig. 1, Table S2). Search for TA proteins revealed that six antitoxins, including GraA (PP_1585), HigA (PP_1198), HigA_2_ (PP_0274), PP_1716, PP_4151 and RelB (PP_2499) and one putative toxin (PP_1717) were detected in wild-type proteome but, as expected, not in the Δ13TA strain. Other antitoxins and toxins were not detectable in either strain. We also analyzed the abundance of proteins encoded by genes located on each side of TA loci to reveal if TA deletions have caused any polar effects on their expression. Most proteins encoded by TA-neighboring genes, which were detectable by mass spectrometry, displayed similar abundances in both wild-type and Δ13TA. Still, we found two proteins, a hypothetical protein encoded by PP_2938 (located downstream of *yefM-yoeB*) and a probable potassium transport protein Kup encoded by PP_1200 (located upstream of *higBA*) that were detected in Δ13TA but not in wild-type. This suggests that deletion of *yefM-yoeB* and *higBA* resulted in upregulation of PP_2938 and Kup, respectively, so that their levels exceeded the detection threshold of the mass spectrometry. Except for these couple of changes, the proteomes of the two strains were very similar, indicating that multiple consecutive rounds of TA deletion procedures have not essentially altered the proteomic profile of Δ13TA strain.

**Figure 1.**
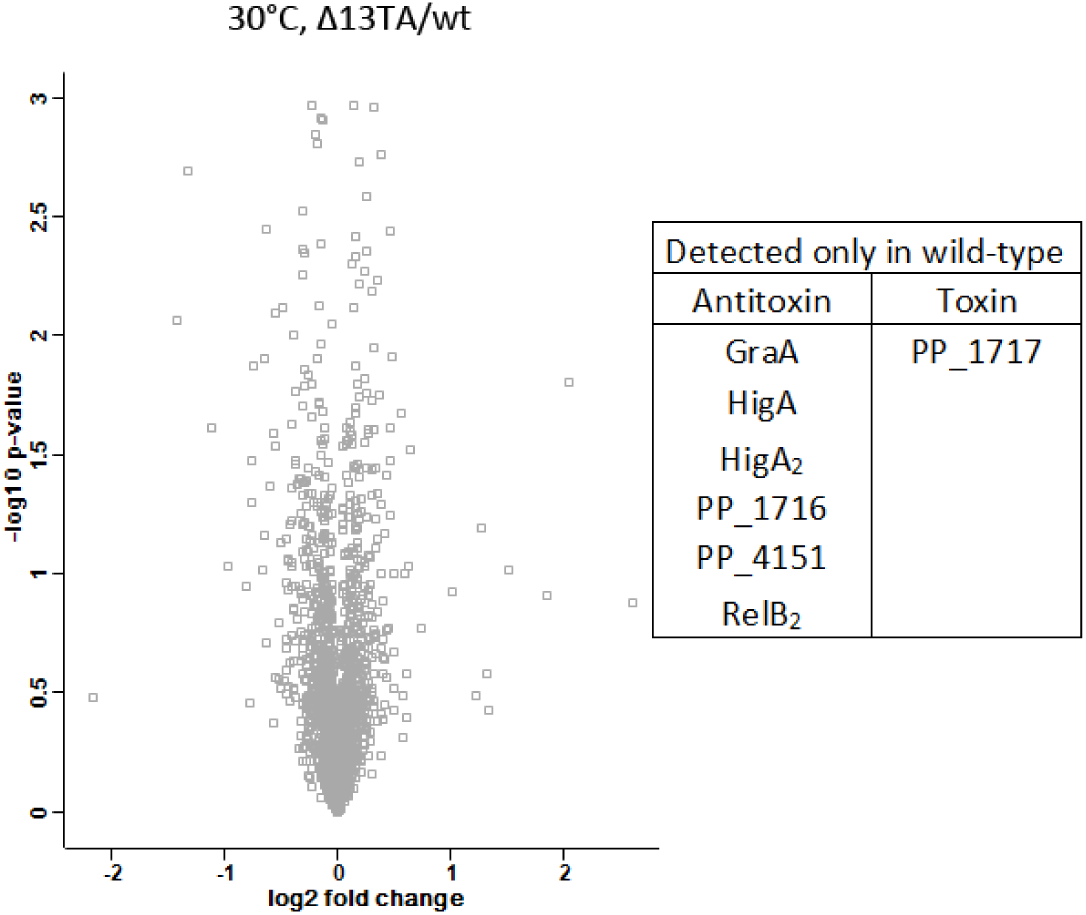
Volcano plot representing the comparison of *P. putida* wild-type and Δ13TA proteome at 30 °C. There were 1867 proteins in the comparison and none of them were significantly differentially expressed between the strains. Six antitoxins and a toxin that were detected only in wild-type proteome are indicated in the table.

### Deletion of 13 TA systems does not affect the stress tolerance, persistence and biofilm formation of *P. putida*

A comparison of the growth of *P. putida* wild-type and Δ13TA strain in rich LB and 2YT as well as in glucose minimal media at 30 °C showed that the absence of multitude TA systems does not affect the overall growth parameters of *P. putida* (Fig. 2A, data for LB is presented only). We next investigated stress tolerance, persistence and biofilm, which are often reported to be influenced by TA systems^14,45^. For analyzing the stress tolerance, the growth of the *P. putida* wild-type and Δ13TA was evaluated on LB medium supplemented with different chemicals that inhibit replication (ciprofloxacin), transcription (rifampicin), translation (tetracycline, kanamycin, streptomycin), or cell wall synthesis (benzylpenicillin) or cause oxidative (4-nitroquinoline 1-oxide, paraquat) or osmotic stress (NaCl). Our data revealed no difference between the stress tolerance of wild-type and Δ13TA strain (Fig. 2B). In order to analyze antibiotic persistence, the two strains were exposed to killing concentrations of streptomycin and benzylpenicillin. Given that the killing curves of *P. putida* wild-type and Δ13TA strain were very similar in both treatments (Fig. 2C and D), the 13 chromosomal TA loci seem not to contribute to antibiotic persistence. Next, the 24-hour biofilm of *P. putida* wild-type and Δ13TA strain grown in LB medium was assayed. As we recorded no differences between the strains (Fig. 2E), the 13 TA systems appear also not to influence the biofilm formation. Thus, under the conditions we used, the absence of 13 chromosomal TA loci does not affect the growth, stress tolerance, persistence or biofilm formation of *P. putida*. These results are actually in line with previous findings that TA loci can be deleted from bacterial chromosome without any clear effects on bacterial behavior^16,27^. Given also the highly similar proteomic profiles of wild-type and Δ13TA strains, the TA deletions seem not to significantly influence the physiology of *P. putida*.

**Figure 2.**
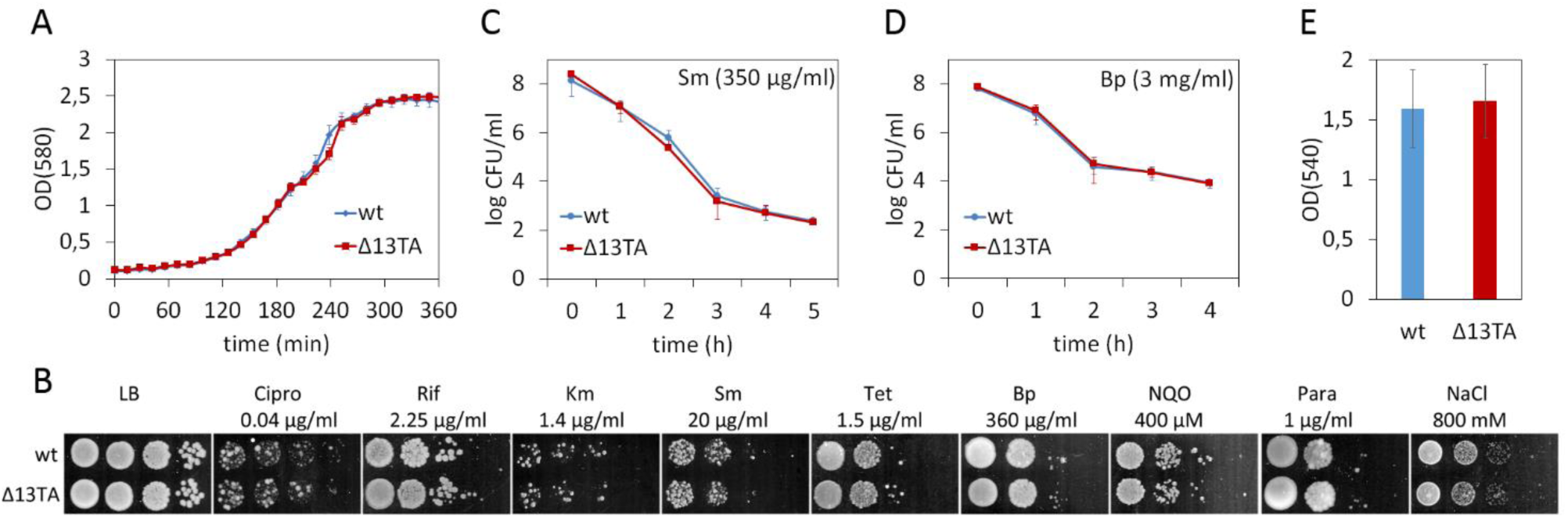
13 TA systems do not affect growth, stress tolerance, persistence and biofilm formation of *P. putida*. (A) Growth curves of *P. putida* wild-type and Δ13TA in LB medium at 30 °C. The strains were grown on microtiter plates. Means from 11 parallels of one measurement with 95% confidence intervals are presented. (B) Stress tolerance plate assays of *P. putida* wild-type and Δ13TA. *P. putida* wild-type and Δ13TA were grown on LB medium for 24 h and on LB medium containing ciprofloxacin (Cipro), rifampicin (Rif), kanamycin (Km), streptomycin (Sm), tetracycline (Tet), benzylpenicillin (Bp), nitroquinoline (NQO), paraquat (Para) or NaCl for 46-48 hours. Final concentrations of the chemicals are indicated. Approximately 5000, 500, 50 and 5 cells were inoculated per spot. (C and D) Killing curves of streptomycin (Sm; C) and benzylpenicillin (Bp; D) of *P. putida* wild-type and Δ13TA. Exponentially growing cultures were treated with Sm (350 µg/ml) or Bp (3 mg/ml) at 30 °C for 5 h (C) or 4 h (D). The surviving cells were determined by CFU plating. Means from four biological replicates with 95% confidence intervals are presented. (E) Biofilm formation of *P. putida* wild-type and Δ13TA. Cells were grown in LB medium at 30 °C for 24 h. Data (means with 95% confidence intervals) from three independent experiments are presented.

### 13 TA systems decrease the competitive fitness of *P. putida* in rich growth conditions but not under stress conditions

To test whether multiple TA loci can influence the competitive fitness of *P. putida* we conducted long-term co-cultivation experiments with 1:1 mixture of the wild-type and Δ13TA strain. We hypothesized that if the TA loci are beneficial then wild-type bacteria will out-compete the Δ13TA strain. If, however, the TA loci are costly, the Δ13TA strain will have the competitive advantage over the wild-type. We marked *P. putida* wild-type and Δ13TA strains with kanamycin and streptomycin resistance genes. In order to recognize the potential effects of resistance marker genes, both antibiotic resistant derivatives of wild-type and Δ13TA strains were constructed. The 1:1 mixtures of wtKm:Δ13TASm as well as wtSm:Δ13TAKm were assayed in parallel experiments during 20-24 days in LB medium at 30 °C. After every two days, the co-cultures were diluted 5000-fold into fresh LB medium. The CFU of wild-type and Δ13TA was determined after every four days. Eight independent co-cultivation experiments revealed that the Δ13TA strain has a statistically significant competitive advantage over the wild-type in seven out of eight trials (Fig. 3, three representative experiments are presented). Only in one parallel of one experiment no statistically significant difference between the wild-type and Δ13TA strains was recorded (data not shown). This indicates that multiple TA systems are costly to *P. putida* growing in rich medium. However, the cost incurred by TA loci seems to be fairly low because Δ13TA strain could never out-compete the wild-type. In fact, we could never detect more than a 1000-fold difference in CFU/ml between the two strains (Fig. 3E and F). Furthermore, some experiments revealed only minor differences between the competitive fitness of the two strains (Fig. 3A and B).

**Figure 3.**
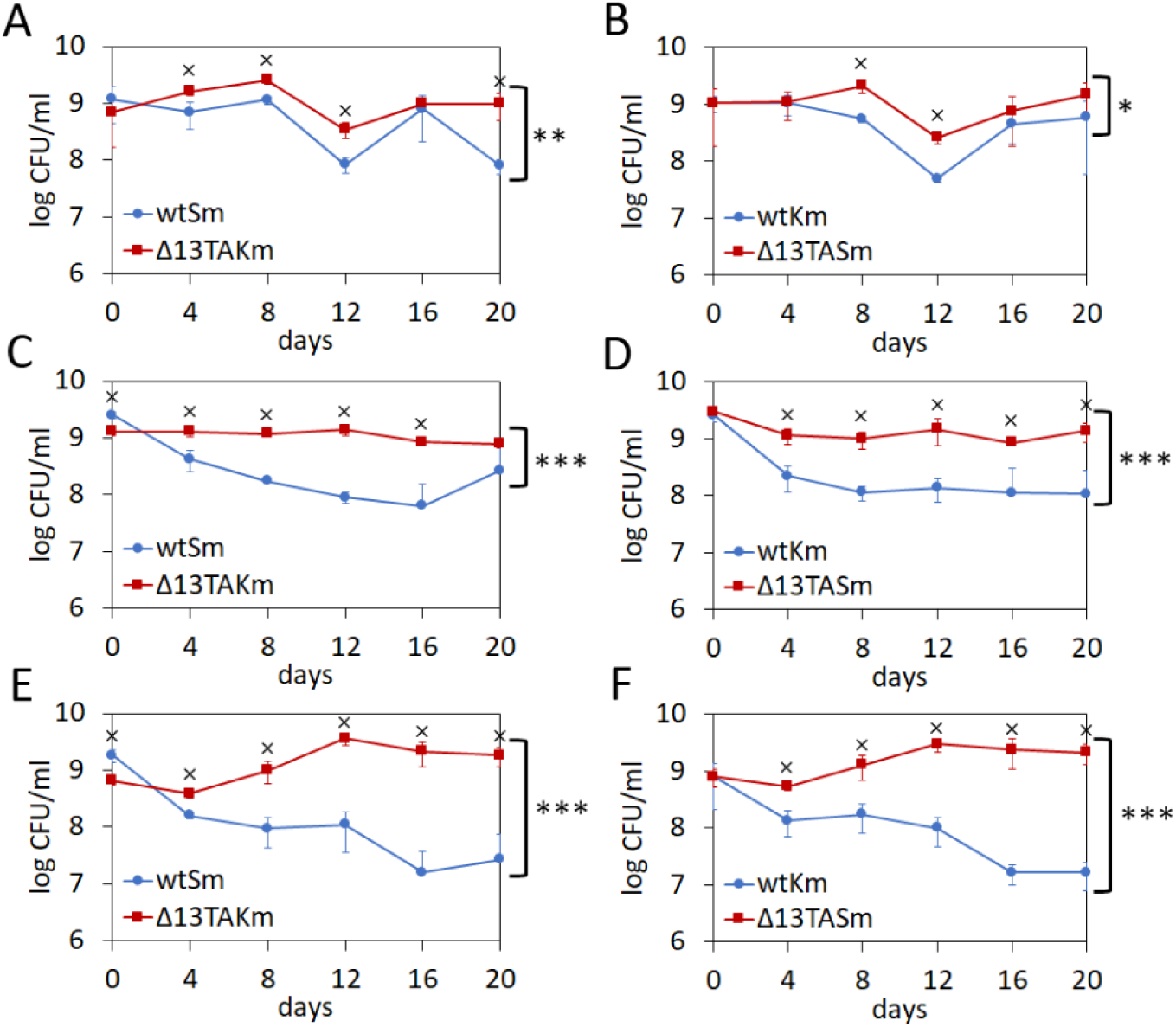
13 TA systems can decrease the competitive fitness of *P. putida* under optimal growth conditions. Co-cultivation of *P. putida* wild-type and Δ13TA strains that are marked with an antibiotic resistance gene (streptomycin or kanamycin). Cells were grown in LB medium at 30 °C, diluted into a fresh LB medium every 2 days and CFU/ml was measured every 4 days. Means from four parallels of one measurement with standard deviation are presented. Two-way ANOVA test was used to evaluate the difference between the two strains over all time points (*, P<0.01; **, P<0.0001; ***, P<0.000001). Wilcoxon rank-sum test was used to evaluate the difference in CFU between the strains in each time point separately (x, P<0.05).

To get more insights into the competitive ability of Δ13TA strain in different conditions, the co-cultivations were performed in different growth media and at suboptimal temperatures as well as under sub-inhibitory concentration of antibiotic stress. Co-cultivation of wild-type and Δ13TA strain in LB medium at 20 °C showed no significant difference between the competitive fitness of the two strains (Fig. 4A and B). Also, neither of the strains could obtain a competitive advantage over the other when grown in M9 minimal medium with glucose (Fig. 4C and D). When sub-inhibitory concentrations of paraquat or ciprofloxacin was added to LB medium we could either not detect a statistically significant difference between wild-type and Δ13TA (Fig. 4E and F, data shown only for paraquat). Yet, we could see that Δ13TA has a slight competitive advantage over wild-type when nitroquinoline or benzylpenicillin was added to LB (Fig. 4G and H, data shown only for nitroquinoline). Notably, we could never detect that wild-type has a competitive advantage over the Δ13TA strain in any of the tested conditions. Finally, we also analysed the competitiveness of the two strains in long-term starvation experiment in M9 minimal medium at 30 °C (Fig. 4I and J). On day zero, stationary phase cells of wtKm and Δ13TASm as well as wtSm and Δ13TAKm were mixed together in 1:1 ratio. The 1:1 mixtures were transferred into fresh M9 minimal medium where they remained during the whole experiment and no additional dilutions to fresh medium were performed. The CFU of wild-type and Δ13TA was determined on days 0, 2, 5, 9, 16, 23, 35 and 50. Again, we could not detect that either of the strains has a competitive advantage over the other (Fig. 4I and J).

**Figure 4.**
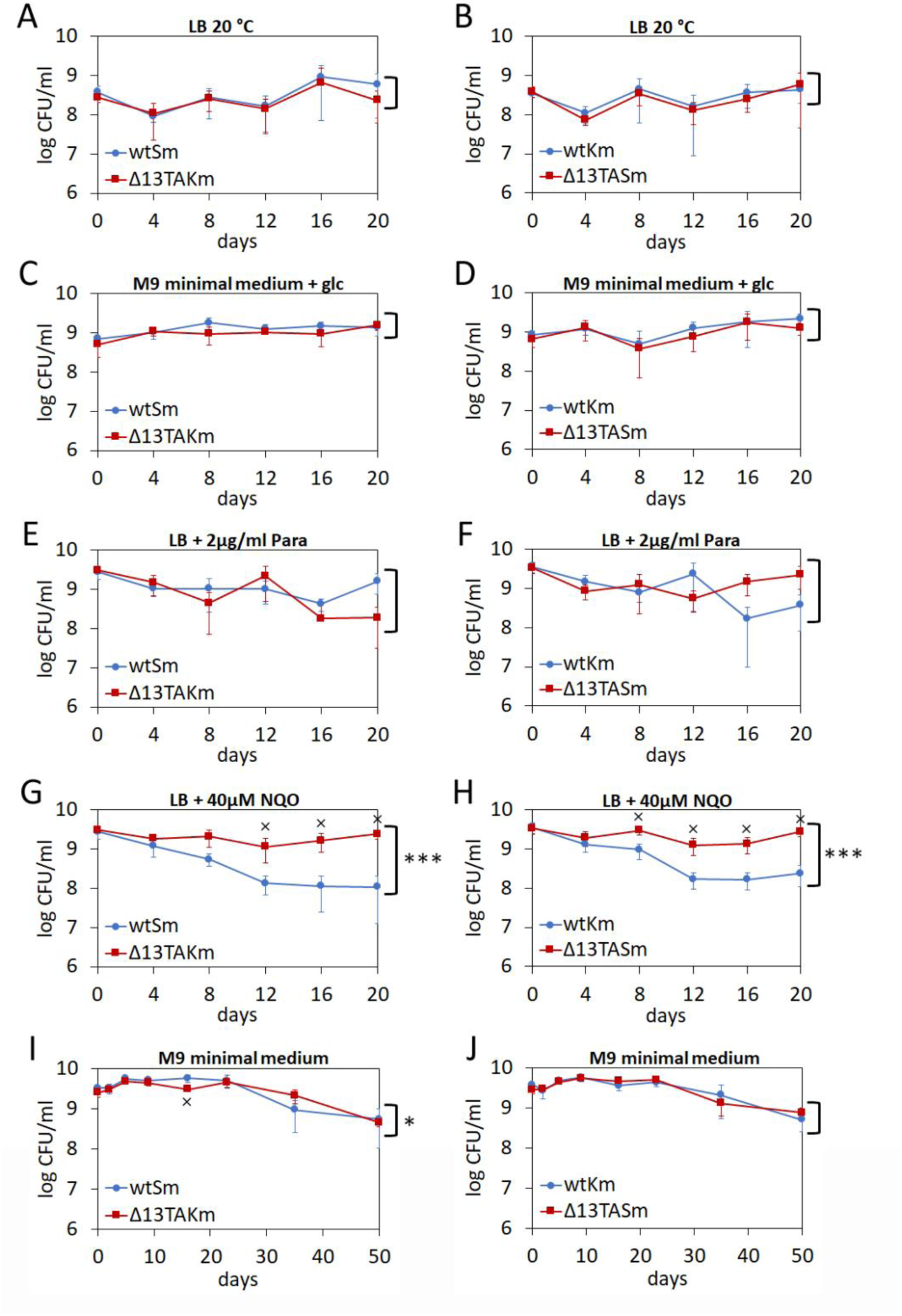
13 TA systems do not contribute to the competitive fitness of *P. putida* under stress conditions. *P. putida* wild-type and Δ13TA strains that are marked with an antibiotic resistance gene (streptomycin or kanamycin) were co-cultivated as 1:1 mixture. (A and B) Bacteria were grown in LB medium at 20 °C. (C and D) Bacteria were grown in M9 minimal medium with glucose at 30 °C. (E to H) Bacteria were grown in LB medium supplemented with paraquat (Para) or nitroquinoline (NQO) at 30 °C. Final concentrations of paraquat and nitroquinoline are indicated. Cells were diluted into a corresponding fresh medium every 2 days and CFU/ml was measured every 4 days (A-H). (I and J) Bacteria were grown in M9 minimal medium at 30 °C. Cells were not diluted to fresh medium during the whole experiment and CFU/ml was measured on days 0, 2, 5, 9, 16, 23, 35 and 50. Except for starvation assay (I and J), at least two independent experiments were performed for each growth condition. Means from at least four parallels of one measurement with standard deviation are presented. Two-way ANOVA test was used to evaluate the difference between the two strains over all time points (*, P<0.01; **, P<0.0001; ***, P<0.000001). Wilcoxon rank-sum test was used to evaluate the difference in CFU between the strains in each time point separately (x, P<0.05).

### TA-encoded proteins exhibit different levels of toxicity

The co-cultivation data indicate that 13 TA loci can carry a fitness cost to *P. putida*. However, the cost seems to be very small and measurable only in some growth conditions (LB medium, LB+nitroquinoline and LB+benzylpenicillin). This raises an interesting question of how toxic the proteins encoded by the chromosomal TA loci actually are. The functionality of TA proteins is usually tested in a kill/rescue assay^46^. Yet, this method involves high copy-number plasmids and artificial overexpression of the toxin that will probably never occur from a single genomic gene copy. Therefore, to study toxins’ effects in their natural genomic context we aimed to construct antitoxin deletion strains from all 13 TA systems. This would evaluate the maximum possible effect the toxin may have to the bacterium. It is expected that antitoxin gene cannot be deleted if the toxin is too noxious and, indeed, deletion of antitoxin gene from five TA loci (*higBA, relE*_*2*_*-higA2*, PP_4151-4152, *yefM-yoeB* and *relB*_*2*_*-parE*) turned out to be impossible (Table 1, indicated in red). At least three independent unsuccessful strain construction trials were performed until we concluded that particular TA system codes for a poisonous toxin. Interestingly, for the other eight TA loci, antitoxin deletion was possible and sequencing verified that the toxin genes had remained intact. It is however important to note that we had to take care that the plasmid, which was used to construct an antitoxin deletion strain would not carry the whole toxin gene because several toxins were deleterious if encoded on the plasmid.

Three TA systems, *graTA, res-xre* and *mqsRA*, encode a toxin that affects the growth rate of the antitoxin deletion strain in optimal growth conditions (LB medium 30 °C) as well as at lower temperature (25 °C) in LB medium and in different stress conditions (Fig. 5). In accordance with previous data, growth inhibition caused by toxin GraT is more severe at lower temperatures^28^. Interestingly, similarly to GraT, the toxic effect of Res also depends on temperature, being more severe at 25 °C (Fig. 5A and B). Differently from that, the growth suppressing effect of toxin MqsR seems not to depend on the temperature (Fig. 5A and B). In compliance with a previous study, we see that GraT can have opposite effects to the stress tolerance of *P. putida*^28^ *– lack of graA* antitoxin increases tolerance to ciprofloxacin, kanamycin, streptomycin and decreases it to tetracycline, benzylpenicillin, nitroquinoline, paraquat and NaCl (Fig. 5C). In addition to GraT, MqsR is the only toxin that has opposite effects to the stress tolerance of *P. putida*. Akin to GraT, MqsR increases tolerance to kanamycin and streptomycin, while decreases it to tetracycline, benzylpenicillin, nitroquinoline, paraquat and NaCl (Fig. 5C). Still, the effects caused by MqsR tend to be smaller than that of GraT. Differently from GraT and MqsR, Res increases sensitivity to kanamycin and streptomycin as well as benzylpenicillin, nitroquinoline, paraquat and NaCl (Fig. 5C).

**Figure 5.**
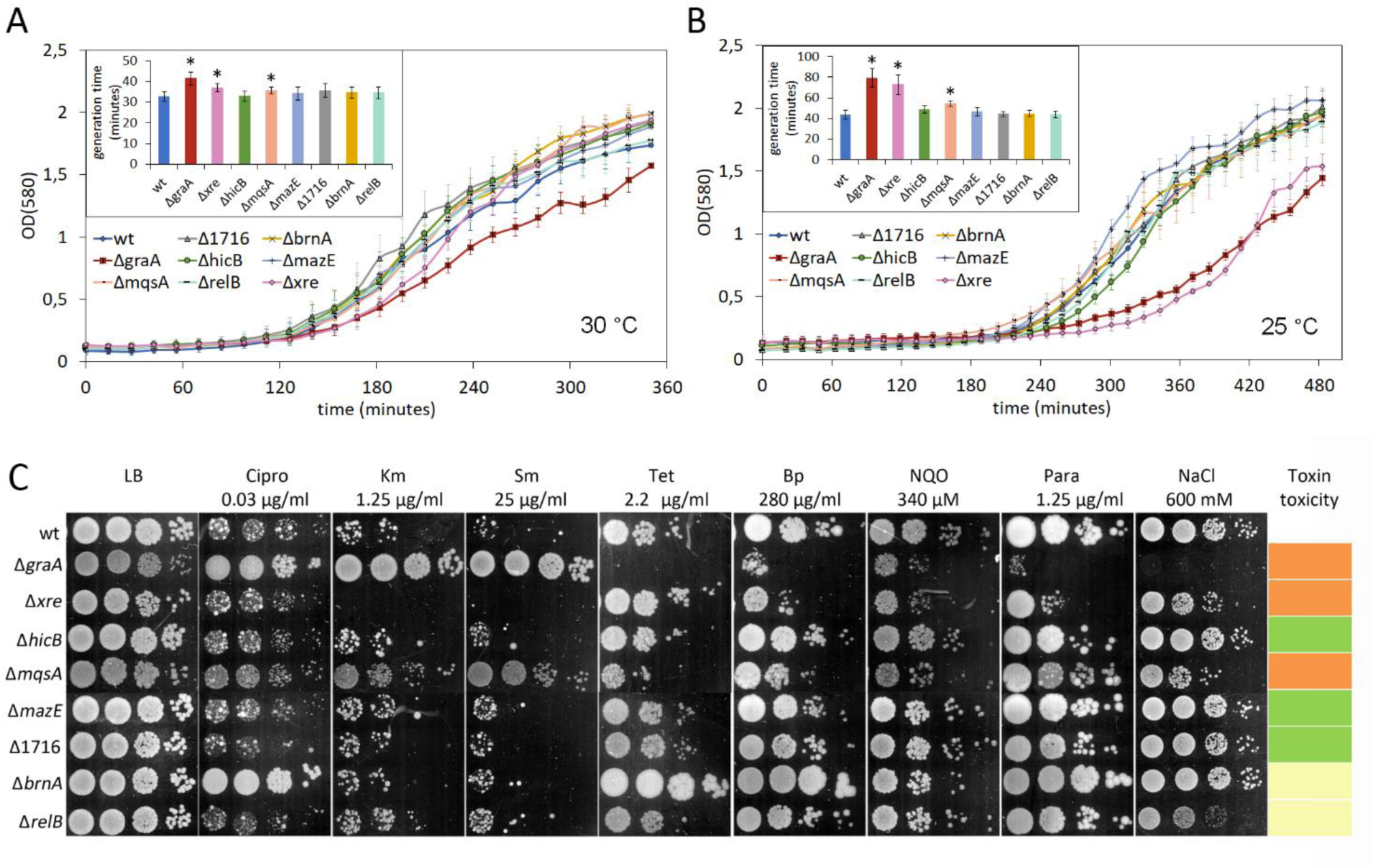
*P. putida* TA systems encode toxins with different toxicity levels. (A and B) Growth curves and generation times of *P. putida* wild-type and antitoxin deletion strains Δ*graA*, Δ*xre*, Δ*hicB*, Δ*mqsA*, Δ*mazE*, Δ1716, Δ*brnA* and Δ*relB* in LB medium at 30 °C (A) and 25 °C (B). The strains were grown on microtiter plates. Data (means with 95% confidence intervals) from at least three independent experiments are presented. (C) Stress tolerance plate assay of *P. putida* wild-type, Δ*graA*, Δ*xre*, Δ*hicB*, Δ*mqsA*, Δ*mazE*, Δ1716, Δ*brnA* and Δ*relB* strains. *P. putida* wild-type and antitoxin deletion strains were grown on LB medium for 24 h and on LB medium containing ciprofloxacin (Cipro), kanamycin (Km), streptomycin (Sm), tetracycline (Tet), benzylpenicillin (Bp), nitroquinoline (NQO), paraquat (Para) or NaCl for 46-48 hours. Final concentrations of the chemicals are indicated. Approximately 5000, 500, 50 and 5 cells were inoculated per spot. Statistically significant changes from wild-type are indicated (*, P<0.05, Student’s T-test). 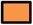- Moderately toxic, affects growth rate and stress tolerance; 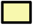- Not toxic, but affects stress tolerance in some conditions; 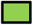- Not toxic in any tested conditions.

Although the toxins of TA systems *brnTA* and *relBE* do not affect the growth rate of *P. putida* in LB medium neither at 30 °C nor 25 °C, they do change the tolerance of *P. putida* to some stress factors (Fig. 5). Toxin BrnT helps *P. putida* manage stress caused by ciprofloxacin, tetracycline, benzylpenicillin and paraquat (Fig. 5C). Out of all the tested conditions toxin RelE only decreases tolerance to NaCl (Fig. 5C). Having such minor effect on *P. putida*’s stress tolerance might indicate that this toxin is losing its toxicity.

For three TA systems, HicAB-1, MazEF and PP_1716-17, we were unable to detect that the deletion of a cognate antitoxin can affect the growth of *P. putida* in any tested conditions indicating that these TA systems do not encode a toxic protein (Fig. 5).

Thus, four types of TA systems were identified in *P. putida* – toxic, moderately toxic and two types of non-toxic systems.

### Diversity and distribution of homologs of 13 *P. putida* TA loci in *Pseudomonads*

The finding that several *P. putida* TA loci are not toxic or incur only mild effects on growth or stress tolerance suggests that these loci have been evolved towards reduced toxicity. Considering that other chromosomal TA systems evolve in similar manner, we hypothesised that disrupted TA loci could be detected among homologs of *P. putida* TA systems. To test this, distribution and diversity of 13 *P. putida* TA loci in genus *Pseudomonas* was analysed by searching 334 fully sequenced *Pseudomonas* strains available in *Pseudomonas* genome database^47^ for putative homologs (Figure S1). *Pseudomonas mesoacidophila* ATCC 31422 was omitted from the analysis as this bacterium was recently reclassified as a member of *Burkholderia cepacia* complex^48^. We first searched for homologs of *P. putida* KT2440 (isogenic to PaW85) antitoxins and found that their distribution in other *Pseudomonads* highly depends on the particular TA system. For example, RelB of the *relBE* and HigA_2_ of the *relE*_*2*_*-higA*_*2*_ system have only 6 and 8 homologs, respectively, but GraA of the *graTA* locus is ubiquitous and has 180 homologs in different *Pseudomonads* (Table 1, Figure S1). Surprisingly, as many as 332 homologs of PP_1716 were found in 334 *Pseudomonas* strains, casting doubt on whether this locus comprises a TA system. Adjacent to the antitoxin homolog the gene homologous to a particular toxin of *P. putida* TA locus was mostly found. Similar to the analysis of antitoxins, the highest number of toxin homologs were detected for GraT (164 full-length homologs) and PP_1717 (334 homologs). As predicted, several toxin genes were truncated and, furthermore, some antitoxin homologs lacked the neighboring toxin gene entirely (Table 1, Figure S1). Toxin gene loss or disruption was most frequent among *xre-res, higBA* and *brnTA* systems where 52%, 27% and 23% of homologous loci, respectively, lacked the intact ORF of the toxin (Table 1). This suggests that these TA loci have been subjected to strong purifying selection, likely to reduce the noxious effects of the toxins.

TA loci are frequently associated with mobile elements that facilitate their horizontal spread^46,49^. Our analysis provides supporting evidence for lateral transfer of TA loci. The best indication of that is the sporadic occurrence of TA homologs on the phylogenetic tree of *Pseudomonads* suggesting that TA loci were acquired through independent insertion events. Independent propagation of TA homologs in different pseudomonads was also supported by analysis of the chromosomal context of TA homologs which revealed that in most cases only the members of *P. putida* clade contain TA homologs in the same chromosomal location as *P. putida* PaW85 (Figure S1). The clear exceptions were, however, homologs of PP_1716-1717 and *relB*_*2*_*-parE* which were found in conserved genomic context across different branches of *Pseudomonads* phylogeny. Remarkably, PP_1716-1717 homologs were located in one genomic context in all *Pseudomonads* they were found. This suggests that rather than being a TA module, the PP_1716-1717 genes most probably belong to the core genome of *Pseudomonads*. Differently from that, the *relB*_*2*_*-parE* module is likely part of a mobile DNA because downstream of this system the integrase gene is located. Furthermore, the integrase as well as several other bordering genes of the *relB*_*2*_*-parE* locus are 100% identical across distantly related *P. putida, P. plecoglossicida NyZ12* and *P. aeruginosa* strains CCUG 70744, AR442, Pa1242, DH01 and PPF-1 suggesting that the DNA region containing *relB*_*2*_*-parE* genes has been gained horizontally.

The other indication of the horizontal spread of TA loci is the presence of several copies of TA homologs in some *Pseudomonads*. For instance, two 100% identical *xre*-related genes were found in different chromosomal position in *P. syringae pv syringae* B301D, *P. azotoformans* S4 and *P. koreensis* D26 (Figure S1). Furthermore, four *xre*-related genes were found in *P. koreensis* P19E3. *P. koreensis* P19E3 possesses one chromosome and three large plasmids^50^, and the four *xre* homologs are distributed between these replicons: two homologs locate in different places in chromosome and other two copies in plasmid 2 and plasmid 3. Interestingly, one chromosomal *xre* homolog (PkP19E3_RS29130) is 100% identical with the plasmid 3-located gene (PkP19E3_RS35085), suggesting gene transfer between replicons.

Thus, phylogenetic analysis of *P. putida* TA system homologs suggests that horizontal gene transfer contributes to spread of TA loci. Finding of incomplete TA loci homologs implies that chromosomal TA loci are under selection pressure and evolve towards less toxic variants.

## Discussion

The idea of chromosomal TA systems acting as stress response elements and contributing to bacterial stress tolerance has been prevalent in TA-related studies. This mainly relies on the ability of toxins to suppress bacterial growth when they are either overexpressed or when chromosomal antitoxin gene is deleted. Non-growing bacteria are known to be protected from environmental insults^51-53^ even if they do not actively combat with stress, as appears to be the case, for example, with most antibiotic-tolerant persister cells^54^. However, despite the potential of TA-encoded toxins to protect bacteria, the deletion of whole TA loci does not usually affect the bacterial fitness^20,27-30,55^, leaving the question of TA systems’ importance in stress tolerance still open.

Data in this study do not support the hypothesis that chromosomal TA systems are important elements in stress tolerance, at least when *P. putida* grows under controlled laboratory conditions. Deletion of 13 chromosomal TA systems had almost no influence on the behaviour of *P. putida*. First, the growth parameters of Δ13TA strain resembled that of wild-type, which was in good agreement with unaltered proteomic profile of the deletion strain. Second, phenotypes that have been usually associated with TA systems, i.e. tolerance to different stress factors, abundance of persister cells and biofilm formation, were also not affected by lack of TA loci. The only effect we could measure was the increased competitive fitness of Δ13TA strain compared to wild-type (Fig. 3). Thus, whereas we could not detect any beneficial effect of TA loci, we show that TA systems confer a fitness cost to *P. putida*. Yet, it should be emphasised that the cost of possessing chromosomal TA loci is quite small and is primarily detected under good growth conditions. If the Δ13TA strain and the wild-type were co-cultivated under sub-optimal or antibiotic stress conditions, the difference in the competitive fitness between the two strains was less noticeable or even undetectable (Fig. 4 A-H). Also, no TA-related fitness costs could be measured when the mix of *P. putida* wild-type and Δ13TA was starved for 50 days (Fig. 4 I and J). Thus, our data suggest that TA loci can impose a slight burden for quickly growing *P. putida* but do not significantly decrease the fitness of non-growing or slowly growing bacteria. Considering that in its natural habitats in soil and water *P. putida* meets mostly poor growth conditions, it is reasonable to presume that multiple chromosomal TA loci do not reduce its fitness in the environment.

Given the noxious nature of TA toxins, the finding that chromosomal TA systems can incur fitness cost to bacterium is expected. So far, however, the potential fitness costs of chromosomal TA loci have attracted only sporadic attention among scientists. One example of TA systems’ fitness cost was observed in *Salmonella* when the deletion of a TA-encoding locus *shpAB* increased the competitiveness of the bacterium in the absence of antibiotic^56^. Yet, TA systems might not always incur fitness costs, despite they encode functional toxins. For example, in a 7 day long co-cultivation assay neither *E. coli* wild-type nor its derivative devoid of five TA systems obtained a competitive advantage over the other^27^. Thus, in accordance with our results, the cost of TA systems seems to be rather small and if the cost is not under direct investigation, it might often remain unnoticed.

The most reasonable explanation for the very low cost of having chromosomal TA systems is that toxins are tightly controlled by their antitoxins. The other reason for low fitness impact of chromosomal TA systems could be that evolution has gradually reduced their toxicity level. In favour to the latter possibility, our data show that more than half of *P. putida*’s toxins (7 out of 12, excluding PP_1716-1717 that is most likely not a TA system after all) are either mildly toxic or have lost their toxicity completely and only five encode for lethal proteins (Table 1). Evidence for evolution reducing toxins’ noxiousness is also supported by the finding of truncated or absent toxin genes among various TA pair homologs both in *Pseudomonads* (Table 1, Figure S1) as well as in *E. coli*^1,57^. *The amount of defective toxin genes among some TA homologs can be rather big, rising up to about 50% and 30% in case of xre-res* system in *Pseudomonads* (Table 1) and *ccd*_*O157*_ system in *E. coli*^57^, *respectively. Such high amount of defective toxin genes among homologs indicates that these TA systems have undergone degeneration during evolution. In this regard, it is interesting to note that the selective pressures acting on either the chromosomal or plasmid-encoded TA systems have been shown to differ significantly: while the plasmid-encoded ccd* systems in *E. coli* are under a strong selective pressure to maintain the toxin function the chromosomal ones are not^57^.

Our bioinformatic analysis further suggests that chromosomal TA systems do not represent a major burden to the bacteria since majority of TA homologs found in different *Pseudomonads* have maintained full toxin ORFs (Table 1). However, we cannot rule out that there are more non-functional or nontoxic TA systems among these seemingly intact TA loci homologs, because bioinformatic analysis cannot detect the impact of different point mutations on TA system functionality. Interestingly, we have observed that inactivating mutations occur in the GraT toxin-encoding gene quickly when antitoxin-deficient Δ*graA* strain is exposed to low temperatures, i.e. under the conditions when GraT strongly suppresses the growth of *P. putida* (unpublished results). Thus, if toxin imposes a real burden to bacterium, it will be quickly neutralized by mutations. One may speculate that due to their noxious nature, the TA systems are subjected to strong selection pressure whenever they enter the genome. However, considering that there are still lots of functional toxins present in bacterial genomes, it might indicate that due to antitoxin’s tight control over the toxin, the genome invading TA systems do not necessary pose an immense threat and, therefore, the selection against them is relatively weak.

Chromosomal TA systems have been most likely acquired from plasmids or bacteriophages via horizontal gene transfer, which is indicated by finding of identical or highly similar TA copies from chromosomes and mobile DNA^1^. Our bioinformatic data also support the idea of plasmids and transposable elements acting as vehicles for spread of TA genes. We often detected transposase or integrase genes in the neighbourhood of TA loci and in some species, e.g. *P. koreensis* P19E3, highly homologous TA operons were found both in its genome and plasmids. Sporadic occurrence of *P. putida* TA systems’ homologs on the phylogenetic tree of *Pseudomonads* provides further evidence that TA genes have been dispersed through independent horizontal transfer events.

The ability of TA systems to propagate horizontally and to stabilize their mobile vehicles may be considered as selfish properties that could be sufficient to ensure the spread and evolutionary success of chromosomal TA systems without them needing to be advantageous to the host bacterium^8,57^. Given that multiple TA loci do not confer any clear benefit to *P. putida*, it is tempting to conclude that they are just selfish genes maintained because of their low fitness cost. Still, we cannot exclude the possibility that *P. putida* TA systems may provide some conditional benefit, i.e. give advantage in certain specific conditions we have not discovered yet. Given that *P. putida* is an environmental bacterium it would be interesting in the future to test the lack of TA systems on phenotypes more related to growth in soil and water.

## Methods

### Bacterial strains, plasmids, and growth conditions

The bacterial strains and plasmids used are listed in Table S3. All strains are derivatives of *P. putida* PaW85^58^, which is isogenic to KT2440^59^. Bacteria were grown in lysogeny broth (LB) or in M9 minimal medium supplemented with 0.2% glucose. If selection was necessary, the growth medium was supplemented with ampicillin (100 µg ml^-1^) or kanamycin (50 µg ml^-1^) for *E. coli* and benzylpenicillin (1500 µg ml^-1^), kanamycin (50 µg ml^-1^) or streptomycin (200 µg ml^-1^) for *P. putida*. Unless noted otherwise, *E. coli* was incubated at 37 °C and *P. putida* at 30 °C. Bacteria were electrotransformed according to the protocol of Sharma and Schimke^60^.

### Construction of plasmids and strains

*P. putida* Δ13TA strain was constructed by sequential deletion of 12 TA loci from *P. putida* Δ*graTA*. For that and for generation of antitoxin deletion strains from *P. putida* PaW85, the pEMG-based plasmids were constructed according to a protocol described elsewhere^61^. The upstream and downstream regions (about 500 bp) of the gene(s) to be deleted were amplified separately and then joined into an approximately 1-kb fragment by overlap extension PCR. Oligonucleotides used in PCR amplification are listed in Table S4. The generated PCR product and pEMG plasmid were digested with restriction enzymes listed in Tabel S5 and then ligated together. Plasmids constructed to generate antitoxin deletion strains carried a part of the toxin gene only (except for pEMG-Δ*hicB-1*) as full-length toxin genes tended to accumulate mutations if encoded on the plasmid. The obtained plasmids were delivered into *P. putida* PaW85 or its deletion derivatives by electroporation, and after 3 h of growth in LB medium the bacteria were plated onto LB agar supplemented with kanamycin. Kanamycin-resistant cointegrates were selected and electrotransformed with the I-SceI expression plasmid pSW(I-SceI). To resolve the cointegrate, the plasmid-encoded I-SceI was induced with 1.5 mM 3-methylbenzoate overnight. Kanamycin-sensitive colonies were selected and the deletions were verified with PCR. The plasmid psW(I-SceI) was eliminated from the deletion strains by growing them overnight in LB medium without antibiotics.

For the construction of wtSm and Δ13TASm or wtKm and Δ13TAKm strains, miniTn7 delivery plasmids pBK-miniTn7-ΩSm or pBK-miniTn7-Km, respectively, together with puXBF13 helper plasmid were coelectroporated into *P. putida* wild-type and Δ13TA strains. Streptomycin or kanamycin resistant bacteria were selected and the miniTn7 insertion to *glmS* locus was verified by PCR.

For construction of pBK-miniTn7-Km, the gentamicin resistance gene, cut from the pBK-miniTn7-ΩGm with SalI and SmaI, was replaced with kanamycin gene. The latter was amplified with oligonucleotide KmSac complementary to both the upstream and downstream sequences of Km^r^ gene in plasmid pUTmini-Tn*5*Km.

### Proteomics

For proteome analysis, bacteria were grown in LB medium overnight, diluted into fresh LB medium for OD_580_ to be 0.1, and grown until mid-exponential phase (OD_580_ ∼1.0). Cells were harvested from three independent cultures per strain. Label-free quantification of whole cell proteomes was performed in the Proteomics Core Facility, Institute of Technology, University of Tartu, Estonia according to previously described protocol^62^. Data analysis was performed with the Perseus software^63^. The whole dataset contained 2,055 different proteins. Parallel samples were grouped together and compared. The analysis included only proteins that were detected in all three parallels (1867 proteins). Mean protein abundances were compared between the two groups using the independent samples Student’s T-test. Benjamini-Hochberg multiple testing correction was applied with the false discovery rate set to 0.05.

### Whole-genome sequencing and bioinformatic analysis

For whole-genome sequencing, genomic DNA of wild-type *Pseudomonas putida* PaW85 and deletion strain Δ13TA was extracted according to the protocol of Thermo Scientific GeneJET Genomic DNA Purification Kit. Both genomes were sequenced using Illumina MiSeq platform with a 100x coverage. Genomes were assembled using SPAdes v3.12.0^64^ and submitted to GenBank (BioProject PRJNA594251). The presence in wild-type and successful deletion of all 13 TA genes in Δ13TA strain was confirmed using homology search. To compare both sequenced genomes to reference strain *Pseudomonas putida* KT2440 (NC_002947.4), sequencing reads were mapped to reference using bowtie2, v2.0.0-beta7^65^ and SNPs and short indels were called using Samtools, v1.9^66^.

### Growth curve and calculation of minimal generation time

To determine growth curves, bacteria were grown overnight in 5 ml of LB medium. The optical densities of bacterial cultures at 580 nm were measured and the bacteria were diluted in LB medium for OD_580_ to be 0.1. Aliquots of 100 µl were transferred into microtiter plate wells and the cells were grown at 30 °C or 25 °C and 400 rpm inside a POLARstar Omega plate reader spectrophotometer. The OD_580_ was measured every 7 min. Data were collected with the Omega data analysis software. Minimal generation time was calculated from the slope of the exponential growth curve according to the formula *G*=*t*/3.3log(*b*/*B*), where *G* is the generation time, *t* is the time interval in minutes, and *B* and *b* are OD_580_ at the beginning and the end of the time interval, respectively.

### Stress tolerance assays

To evaluate stress tolerance, bacteria were grown overnight in 5 ml of LB medium. 10-fold serial dilutions of the cultures were spotted as 5 µl drops onto LB plates supplemented with different chemicals (specified in Results) and incubated at 30 °C for 24 h or 46-48 h.

### Persistence assays

To determine the amount of persisters left after antibiotic treatment, bacteria were grown overnight in 5 ml of LB medium. The overnight grown cultures were diluted 50-fold into LB medium and grown at 30 °C to exponential growth phase (OD_580_ ∼ 0.7-0.9). A 1.4-ml portion of each culture was transferred to an Eppendorf tube and treated with 350 µl/ml streptomycin or 3 mg/ml benzylpenicillin with shaking at 30 °C. CFU/ml was determined before the antibiotic was added and at various time points during the assay. For that, 200 µl of bacterial culture was centrifuged and resuspended in M9 buffer, and 10-fold serial dilutions of the cultures were spotted as 5 µl drops onto LB plates and incubated at 30 °C overnight.

### Biofilm formation assay

The overnight in 5 ml of LB medium grown cultures were diluted 20-fold into LB medium and aliquots of 100 µl were transferred into polystyrene microtiter plate wells. The microtiter plate was incubated at 30 °C for 24 h. Next, 25 µl of 1% crystal violet was added for 15 min to stain the cells. Then the wells were washed twice with 150 µl of H_2_O and after that 180 µl of 96 % ethanol was added twice to the wells to extract crystal violet from cells in biofilm. Finally, 100 µl of crystal violet in ethanol was diluted into 200 µl of water in another multi-well plate and the OD_540_ of the solutions was measured with a microtitre plate reader (Tecan Sunrise-Basic).

### Competition assay

*P. putida* wild-type and Δ13TA strains that are marked with an antibiotic resistance gene (streptomycin or kanamycin) were grown overnight in 5 ml LB medium or M9 minimal medium with glucose at 30 °C. The optical densities of the bacterial cultures at 580 nm were measured and the 1:1 mixture containing the equal amounts of wtSm and Δ13TAKm or wtKm and Δ13TASm cells were prepared. The 1:1 mixture was diluted 5000-10 000-fold (about 10^5^ cells per ml) into fresh 5 ml of LB or M9 minimal medium with glucose and grown at 30 °C. If indicated, the growth temperature was lowered to 20 °C or the medium was supplemented with an antibiotic to induce stress. Cells were diluted into a corresponding fresh medium every 2 days and CFU/ml was measured every 4 days.

For starvation assay, the 1:1 mixtures of wtSm and Δ13TAKm or wtKm and Δ13TASm were prepared from LB-grown overnight cultures as described above. The cells were then centrifuged out from the medium (about 8 ml) and washed two times with 1x M9. The washed mixtures were transferred into 5 ml of M9 minimal medium and grown at 30 °C for 50 days. No additional dilutions to fresh medium were performed. CFU/ml was measured on days 0, 2, 5, 9, 16, 23, 35 and 50.

### Phylogenetic analysis of TA homologs

Search for *P. putida* PaW85 TA systems’ homologs within the *Pseudomonas* Genome Database^47^ was conducted using the DIAMOND BLAST program. The antitoxin and toxin protein sequences of *P. putida* KT2440 (isogenic to strain PaW85) were queried against complete genomes of 334 *Pseudomonads* (September 2019) with default settings, except that 30% identity cutoff was used. If only one protein of the putative TA pair was detected (query with antitoxin gave mostly more homologs than with toxin), then the manual search was conducted to find unannotated or truncated ORFs.

### Phylogenetic tree of *Pseudomonads*

We included 334 available *Pseudomonads* with genome sequencing status marked as complete in *Pseudomonas* Genome Database^47^. Concatenated *rpoB, rpoD, gyrB* and *guaA* gene sequences from each genome were aligned using MUSCLE, v3.8.31^67^. Phylogenetic tree was calculated with FastTree, v2.1.4 SSE3^68^ and visualized and edited in iTOL^69^.

## Data availability

Whole sequences of *P. putida* PaW85 and Δ13TA are submitted to GenBank (BioProject PRJNA594251).

## Acknowledgements

This work was supported by the Estonian Research Council grants PUT1351 and IUT34-11 and by the EU European Regional Development Fund grant No. 2014-2020.4.01.15-0012.

## Author Contributions

R.H. conceived the project, S.R. and H.T. performed the experiments, S.R., A.B. and M.R. performed data analysis, S.R. and R.H. wrote the manuscript, all authors reviewed and approved the manuscript.

