## Supplementary Figure 1 for "*Pseudomonas putida* chromosomal toxin-antitoxin systems carry neither clear fitness benefits nor big costs"

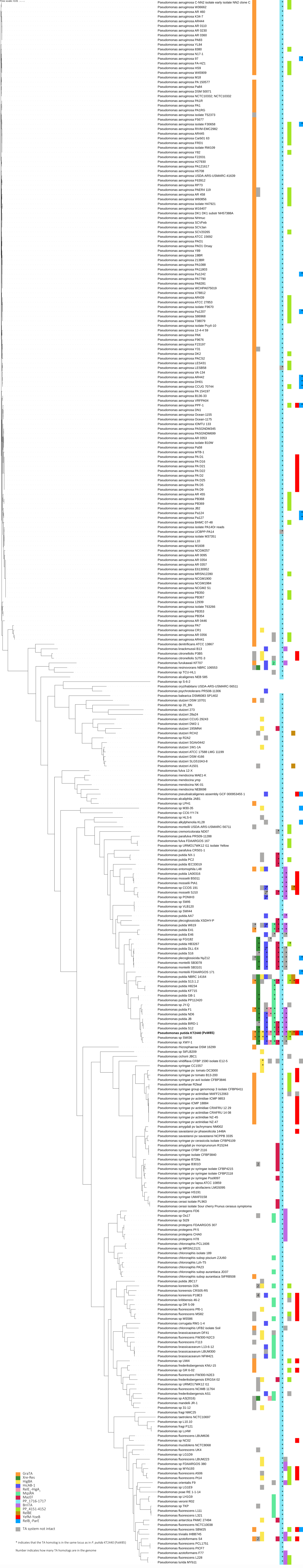


**Supplementary Figure 1.** Distribution and diversity of *P. putida* TA systems' homologs among 334 fully sequenced *Pseudomonas* strains.
