## Supplementary Tables 1 & 3-5 for "*Pseudomonas putida* chromosomal toxin-antitoxin systems carry neither clear fitness benefits nor big costs"

**Supplementary Table 1**. Mutational differences between *P. putida* KT2440, *P. putida* PaW85 and the PaW85 derivative devoid of 13 TA systems.

| Position in reference strain KT2440 | KT2440 (NC_002947.4) | PaW85[a] | Δ13TA[a] | Locus_tag | Annotation in KT2440 reference genome |
|---|---|---|---|---|---|
| 278742 | T | TA | TA | | intergenic region |
| 307865 | G | GC | GC | PP_0253 | GeneID:1043829, pseudogene, fragment of phosphoenolpyruvate carboxykinase (ATP) |
| 336124 | A | . | AT | PP_0278 | GeneID:1043886, hypothetical protein |
| 353067 | A | G | G | | intergenic region |
| 499203 | A | G | G | | intergenic region |
| 1070246 | T | . | TGA | | intergenic region |
| 1126645 | A | AC | AC | | intergenic region |
| 1419548 | G | GC | . | | intergenic region |
| 1499480 | C | A | A | | intergenic region |
| 1499481 | A | C | C | | intergenic region |
| 1499498 | T | G | G | | intergenic region |
| 1916247 | C | . | T | PP_1715 | GeneID:1043474, hypothetical protein |
| 1916345 | G | . | A | | intergenic region |
| 1916352 | A | . | G | | intergenic region |
| 1932638 | G | GC | GC | | intergenic region |
| 2069519 | T | C | . | | intergenic region |
| 2069526 | T | C | . | | intergenic region |
| 3072261 | C | T | T | PP_2682 | GeneID:1046138, yiaY, Fe-containing alcohol dehydrogenase |
| 3343032 | G | . | C | | intergenic region |
| 3399678 | G | C | C | PP_3005 | GeneID:1043105, hypothetical protein |
| 3523296 | C | . | A | PP_3114 | GeneID:1043961, transposase |
| 3523297 | A | . | G | PP_3114 | GeneID:1043961, transposase |
| 3523305 | A | T | . | PP_3114 | GeneID:1043961, transposase |
| 3523311 | C | T | . | PP_3114 | GeneID:1043961, transposase |
| 3845703 | C | CG | CG | | intergenic region |
| 3951159 | T | C | C | | intergenic region |
| 3951161 | A | T | T | | intergenic region |
| 4068175 | C | . | G | PP_3583 | GeneID:1046361, mdtC, multidrug transporter membrane protein |
| 4536662 | G | C | C | | intergenic region |
| 4536664 | C | G | G | | intergenic region |
| 4536674 | A | C | C | PP_4025 | GeneID:1042069, transposase |
| 4536675 | G | A | A | PP_4025 | GeneID:1042069, transposase |
| 4536683 | T | A | A | PP_4025 | GeneID:1042069, transposase |
| 4536689 | T | C | C | PP_4025 | GeneID:1042069, transposase |
| 4586030 | C | T | T | | intergenic region |
| 4586031 | T | C | C | | intergenic region |
| 4740804 | CT | C | C | | intergenic region |
| 4740816 | T | G | G | | intergenic region |

| | | | | | |
|---|---|---|---|---|---|
| 4740819 | T | C | C | | intergenic region |
| 4740820 | G | T | T | | intergenic region |
| 4741230 | A | . | G | PP_5662 | pseudogene, GeneID:26969972, unknown function |
| 4741232 | C | . | T | PP_5662 | pseudogene, GeneID:26969972, unknown function |
| 4741235 | C | . | T | PP_5662 | pseudogene, GeneID:26969972, unknown function |
| 4741238 | C | . | G | PP_5662 | pseudogene, GeneID:26969972, unknown function |
| 4741240 | A | . | C | PP_5662 | pseudogene, GeneID:26969972, unknown function |
| 4741241 | C | . | T | PP_5662 | pseudogene, GeneID:26969972, unknown function |
| 4741258 | A | G | G | PP_5662 | pseudogene, GeneID:26969972, unknown function |
| 4741263 | A | C | C | | intergenic region |
| 4741272 | A | G | G | | intergenic region |
| 4741274 | A | C | C | | intergenic region |
| 4741276 | G | C | C | | intergenic region |
| 4741278 | C | T | T | | intergenic region |
| 4741289 | C | G | G | | intergenic region |
| 4741292 | C | T | T | | intergenic region |
| 4741294 | C | G | G | | intergenic region |
| 4741296 | A | C | C | | intergenic region |
| 4741298 | C | T | T | | intergenic region |
| 4980585 | G | GGGC | GGGC | | intergenic region |
| 5182339 | G | . | A | | intergenic region |
| 5311566 | T | A | . | | GeneID:1044736, 16S ribosomal RNA |
| 5555608 | G | GGCC | GGCC | | intergenic region |
| 5674750 | C | CG | CG | | intergenic region |
| 5674753 | G | C | C | | intergenic region |
| 5681415 | C | CCGGG | . | | intergenic region |
| 6013885 | T | C | C | | intergenic region |

[a] . Indicates identical nucleotide to reference *P. putida* KT2440

**Supplementary Table 3.** Strains and plasmids

| Strain or plasmid | Genotype or characteristic(s) | Source or reference |
|---|---|---|
| *E. coli* strain | | |
| DH5α λ*pir* | λpir lysogen of DH5α | (1) |
| *P. putida* strains | | |
| PaW85 | Wild-type, isogenic to KT2440 | (2) |
| Δ*graTA* | PaW85 Δ*graTA* | (3) |
| Δ2TA | Δ*graTA* Δ*res-xre* | this study |
| Δ3TA | Δ2TA Δ*higBA* | this study |
| Δ4TA | Δ3TA Δ*hicAB-1* | this study |
| Δ5TA | Δ4TA Δ*relE*$_2$-*higA*$_2$ | this study |
| Δ6TA | Δ5TA Δ*mqsRA* | this study |
| Δ7TA | Δ6TA Δ*mazEF* | this study |
| Δ8TA | Δ7TA ΔPP_1716-1717 | this study |
| Δ9TA | Δ8TA Δ*brnTA* | this study |
| Δ10TA | Δ9TA ΔPP_4151-4152 | this study |
| Δ11TA | Δ10TA Δ*relBE* | this study |
| Δ12TA | Δ11TA Δ*yefM-yoeB* | this study |
| Δ13TA | Δ12TA Δ*relB*$_2$-*parE* | this study |
| wtSm | PaW85 with miniTn7-ΩSm in *glmS* locus | this study |
| wtKm | PaW85 with miniTn7-Km in *glmS* locus | this study |
| Δ13TASm | Δ13TA with miniTn7-ΩSm in *glmS* locus | this study |
| Δ13TAKm | Δ13TA with miniTn7-Km in *glmS* locus | this study |
| Δ*graA* | PaW85 Δ*graA* | (3) |
| Δ*xre* | PaW85 Δ*xre* | this study |
| Δ*hicB* | PaW85 Δ*hicB* | this study |
| Δ*mqsA* | PaW85 Δ*mqsA* | this study |
| Δ*mazE* | PaW85 Δ*mazE* | this study |
| Δ1716 | PaW85 Δ1716 | this study |
| Δ*brnA* | PaW85 Δ*brnA* | this study |
| Δ*relB* | PaW85 Δ*relB* | this study |
| Plasmids | | |
| pEMG | Plasmid for homologous recombination, *lacZα* with two flanking I-SceI sites ($Km^r$) | (1) |
| pSW(I-SceI) | Plasmid coding for I-SceI endonuclease for allelic exchange experiments ($Bp^r$) | (4) |
| pEMG-Δ*res-xre* | pEMG containing chimeric DNA fragment for deleting *res-xre* ($Km^r$) | this study |
| pEMG-Δ*higBA* | pEMG containing chimeric DNA fragment for deleting *higBA* ($Km^r$) | this study |
| pEMG-Δ*hicAB-1* | pEMG containing chimeric DNA fragment for deleting *hicAB-1* ($Km^r$) | this study |
| pEMG-Δ*relE*$_2$-*higA*$_2$ | pEMG containing chimeric DNA fragment for deleting *relE*$_2$-*higA*$_2$ ($Km^r$) | this study |
| pEMG-Δ*mqsRA* | pEMG containing chimeric DNA fragment for deleting *mqsRA* ($Km^r$) | this study |
| pEMG-Δ*mazEF* | pEMG containing chimeric DNA fragment for deleting mazEF ($Km^r$) | this study |
| pEMG-Δ1716-1717 | pEMG containing chimeric DNA fragment for deleting PP_1716-1717 ($Km^r$) | this study |
| pEMG-Δ*brnTA* | pEMG containing chimeric DNA fragment for deleting brnTA ($Km^r$) | this study |
| pEMG-Δ4151-4152 | pEMG containing chimeric DNA fragment for deleting PP_4151-4152 ($Km^r$) | this study |

| | | |
|---|---|---|
| pEMG-Δ*relBE* | pEMG containing chimeric DNA fragment for deleting *relBE* ($Km^r$) | this study |
| pEMG-Δ*yefM-yoeB* | pEMG containing chimeric DNA fragment for deleting *yefM-yoeB* ($Km^r$) | this study |
| pEMG-Δ*relB₂-parE* | pEMG containing chimeric DNA fragment for deleting *relB₂-parE* ($Km^r$) | this study |
| pEMG-Δ*xre* | pEMG containing chimeric DNA fragment for deleting *xre* ($Km^r$) | this study |
| pEMG-Δ*higA* | pEMG containing chimeric DNA fragment for deleting *higA* ($Km^r$) | this study |
| pEMG-Δ*hicB-1* | pEMG containing chimeric DNA fragment for deleting *hicB-1* ($Km^r$) | this study |
| pEMG-Δ*higA₂* | pEMG containing chimeric DNA fragment for deleting *higA₂* ($Km^r$) | this study |
| pEMG-Δ*mqsA* | pEMG containing chimeric DNA fragment for deleting *mqsA* ($Km^r$) | this study |
| pEMG-Δ*mazE* | pEMG containing chimeric DNA fragment for deleting *mazE* ($Km^r$) | this study |
| pEMG-Δ*1716* | pEMG containing chimeric DNA fragment for deleting PP_1716 ($Km^r$) | this study |
| pEMG-Δ*brnA* | pEMG containing chimeric DNA fragment for deleting *brnA* ($Km^r$) | this study |
| pEMG-Δ*4151* | pEMG containing chimeric DNA fragment for deleting PP_4151 ($Km^r$) | this study |
| pEMG-Δ*relB* | pEMG containing chimeric DNA fragment for deleting *relB* ($Km^r$) | this study |
| pEMG-Δ*yefM* | pEMG containing chimeric DNA fragment for deleting *yefM* ($Km^r$) | this study |
| pEMG-Δ*relB₂* | pEMG containing chimeric DNA fragment for deleting *relB₂* ($Km^r$) | this study |
| puXBF13 | Plasmid coding for the Tn7 transposition proteins ($Amp^r$ *mob*$^+$) | (5) |
| pBK-miniTn7-ΩSm | pUC19-based delivery plasmid for miniTn7-ΩSm ($Amp^r$ $Sm^r$) | (6) |
| pBK-miniTn7-ΩGm | pUC19-based delivery plasmid for miniTn7-ΩGm ($Amp^r$ $Gm^r$) | (6) |
| pBK-miniTn7-Km | pBK-miniTn7-ΩGm derivative, Gm is replaced with Km gene ($Amp^r$ $Km^r$) | this study |

**Supplementary Table 4.** Oligonucleotides

| Name | Sequence (5'-3')[a] | Use |
|---|---|---|
| 2433Sac | gagagctcgcacgtgattgtgttct | construction of pEMG-Δxre-res and pEMG-Δxre |
| 2433pikk | ctgtggtgaccacgcccta-gcggtatatgccgggtcat | construction of pEMG-Δxre-res |
| 2434taga | tagggcgtggtcaccacag | construction of pEMG-Δxre-res |
| 2434Xba | ggtctagatgtctttttcggcgcaca | construction of pEMG-Δxre-res and pEMG-Δxre |
| 1199BHI | aggatccccagcacgccat | construction of pEMG-ΔhigBA and pEMG-ΔhigA |
| 1199ees | ggcgctcatctaagttgtac | construction of pEMG-ΔhigBA |
| 1198pikk | gtacaacttagatgagcgcc-gagctaagtacacgaaagc | construction of pEMG-ΔhigBA |
| 1198Sac | tggagctcacggatgctgccttctt | construction of pEMG-ΔhigBA and pEMG-ΔhigA |
| 1479Eco | gtgaattcgagaatcgaatccgcct | construction of pEMG-ΔhicAB-1 and pEMG-ΔhicB |
| 1479pikk | ccaacaatcagctggtatcatgcagtagcttaatcccgc | construction of pEMG-ΔhicAB-1 |
| 1480ees | tgataccagctgattgttgg | construction of pEMG-ΔhicAB-1 |
| 1480Bam | aaggatccaacgactccaactacgg | construction of pEMG-ΔhicAB-1 and pEMG-ΔhicB |
| 274Sal | ctggtcgacaagcactactacagc | construction of pEMG-ΔrelE$_2$-higA$_2$ and pEMG-ΔhigA$_2$ |
| 274pikk | cggctatcgagcctgtcaactccgatccagatatctgtc | construction of pEMG-ΔrelE$_2$-higA$_2$ |
| 274stop | ttgacaggctcgatagccg | construction of pEMG-ΔrelE$_2$-higA$_2$ and pEMG-ΔhigA$_2$ |
| 275Sac | atgagctcaatcgcacccg | construction of pEMG-ΔrelE$_2$-higA$_2$ and pEMG-ΔhigA$_2$ |
| 4204Acc | tgggtaccagaactggttcggtgg | construction of pEMG-ΔmqsRA and pEMG-ΔmqsA |
| 4204pikk | gccaaatttaacctggaaggctgcctgataatggcagcc | construction of pEMG-ΔmqsRA |
| 4205ees | ccttccaggttaaatttggc | construction of pEMG-ΔmqsRA |
| 4205(Sal) | tcgtcttcctcaacattggc | construction of pEMG-ΔmqsRA and pEMG-ΔmqsA |
| 769Hind | gctaagcttgtacaaactgggtgtg | construction of pEMG-ΔmazEF and pEMG-ΔmazE |
| 770ees | gagtatctcccaaggtagat | construction of pEMG-ΔmazEF |
| 771pikk | tctaccttgggagatactcaaccggttcccttattcaca | construction of pEMG-ΔmazEF |
| 772BHI | acggatccttgatggaacgcacgat | construction of pEMG-ΔmazEF and pEMG-ΔmazE |
| 1716Sac | aagagctcggccaactgcgtgaag | construction of pEMG-Δ1716-1717 and pEMG-Δ1716 |
| 1716TAdelpikk | ggaatgggcaacgcttcagcagcatgtaatccgtaggca | construction of pEMG-Δ1716-1717 |
| 1717stop | ctgaagcgttgcccattcc | construction of pEMG-Δ1716-1717 |
| 1718Bam | ttggatcctggtgcagggctatttg | construction of pEMG-Δ1716-1717 and pEMG-Δ1716 |
| 4528Xba | cgtctagaagctaccgattatcggg | construction of pEMG-ΔbrnTA and pEMG-ΔbrnA |
| 4528pikk | aaccagagcaatatccgcaaccttggcaggtccca | construction of pEMG-ΔbrnTA |
| 4530ees | ttgcggatattgctctggtt | construction of pEMG-ΔbrnTA |
| 4531Sac | ttgagctctgcctgtttcaactcct | construction of pEMG-ΔbrnTA and pEMG-ΔbrnA |
| 4151Eco | acgaattctgctggatgaacccac | construction of pEMG-Δ4151-4152 and pEMG-Δ4151 |
| 4151-52delpikk | ggaacttgaatgctgaaagcacatggctagagaggt | construction of pEMG-Δ4151-4152 |
| 4151-52del | tgctttcagcattcaagttcc | construction of pEMG-Δ4151-4152 |
| 4152Kpn | acgggtaccggcaaaccttg | construction of pEMG-Δ4151-4152 and pEMG-Δ4151 |
| 1266Sac | aggagctcaagcgtacgcacatcg | construction of pEMG-ΔrelBE and pEMG-ΔrelB |
| 1267TAdelpikk | catcaccgtggtaatgcacagatgatgcgggaggc | construction of pEMG-ΔrelBE |
| 1268start | gtgcattaccacggtgatg | construction of pEMG-ΔrelBE |
| 1269Bam | agggatcctctgccaaggattccaa | construction of pEMG-ΔrelBE and pEMG-ΔrelB |
| 2938Sac | tagagctccaatttacggtgacagcg | construction of pEMG-ΔyefM-yoeB |
| 2939TAdelpikk | gtccaaagagcaacccttacgcggtccttttgtaacgg | construction of pEMG-ΔyefM-yoeB |
| 2940start | gtaagggttgctctttggac | construction of pEMG-ΔyefM-yoeB |
| 2940Xba | attctagagacagaatacaccgctg | construction of pEMG-ΔyefM-yoeB and pEMG-ΔyefM |

| | | |
|---|---|---|
| 2498EcoRI | tagaattccagattcgtcctgcaac | construction of pEMG-ΔrelB$_2$-parE and pEMG-ΔrelB$_2$ |
| 2499TAdelpikk | ctgtccagtctcagtagcccccgttatctaaagg | construction of pEMG-ΔrelB$_2$-parE |
| 2500stop | ggctactgagactggacag | construction of pEMG-ΔrelB$_2$-parE |
| 2501Xba | catctagaaaggcacctggttcatc | construction of pEMG-ΔrelB$_2$-parE and pEMG-ΔrelB$_2$ |
| 2433_A_pikk | caggccatagccctcggcatttgccgacaagta | construction of pEMG-Δxre |
| 2433_A_del | tgccgagggctatggcctg | construction of pEMG-Δxre |
| higApikk | aaagggtggaaataatg-gaaagccaccaggccctag | construction of pEMG-ΔhigA |
| higAdel | tttccattatttccacccttt | construction of pEMG-ΔhigA |
| hicB-1pikk | tgacctaccgcggctatgcagtagcttaatcccgc | construction of pEMG-ΔhicB |
| hicB-1del | gcatagccgcggtaggtca | construction of pEMG-ΔhicB |
| 274_pikk_2 | cggctatcgagcctgtcaa-tcgaattgaaatatcgcgcg | construction of pEMG-ΔhigA$_2$ |
| 4204ees | ctcatggttacaactccttg | construction of pEMG-ΔmqsA |
| del4204 | caaggagttgtaaccatgaggaagtgcgtactgcc | construction of pEMG-ΔmqsA |
| 770ees | gagtatctcccaaggtagat | construction of pEMG-ΔmazE |
| 771pikk | tctaccttgggagatactc-aaccggttcccttattcaca | construction of pEMG-ΔmazE |
| 1716stop | ctggctggagcgagccatc | construction of pEMG-Δ1716 |
| 1716delpikk | gatggctcgctccagccagcatgtaatccgtaggca | construction of pEMG-Δ1717 |
| 4530stop | gatatccttgtcgtcctgtc | construction of pEMG-ΔbrnA |
| 4530delpikk | gacaggacgacaaggatatcgcaaccttggcaggtccca | construction of pEMG-ΔbrnA |
| 4151lopp | gcagcccaagctcaggaat | construction of pEMG-Δ4151 |
| 4151del_pikk | aattcctgagcttgggctgc-aagcacatggctagagaggt | construction of pEMG-Δ4151 |
| 1268lopp | cgacctggatcaacctctgt | construction of pEMG-ΔrelB |
| 1268del_pikk | acagaggttgatccaggtcgcgattgcattgccttgtca | construction of pEMG-ΔrelB |
| 2940del | ttcggcgcgagcctggcta | construction of pEMG-ΔyefM |
| 2940del_pikk | tagccaggctcgcgcc-gaaccagaagcagcggaaca | construction of pEMG-ΔyefM |
| 2939Sac | tagagctcgaagaagtagaccaggc | construction of pEMG-ΔyefM |
| 2499del | gctgaggcagagttacccaa | construction of pEMG-ΔrelB$_2$ |
| 2499del_pikk | ttgggtaactctgcctc-agcccccgttatctaaagg | construction of pEMG-ΔrelB$_2$ |
| Tn7R109 | cagcataactggactgatttcag | verification of miniTn7 insertion to *glmS* locus |
| Tn7GlmS | aatctggccaagtcggtgac | verification of miniTn7 insertion to *glmS* locus |
| KmSac | caggagctcgttcgatttattcaacaaagcc | amplification of Km gene |

[a] The sites of restriction enzymes used in cloning are underlined.

**Supplementary Table 5**. Construction strategy of pEMG-based plasmids for gene deletions

| **Plasmid** | **Restriction enzymes used to digest PCR product**[a] | **Restriction enzymes used to digest pEMG plasmid**[a] |
|---|---|---|
| pEMG-Δ*res-xre* | SacI and XbaI | SacI and XbaI |
| pEMG-Δ*higBA* | BamHI and SacI | BamHI and SacI |
| pEMG-Δ*hicAB-1* | BamHI and EcoRI | BamHI and EcoRI |
| pEMG-Δ*relE*$_2$-*higA*$_2$ | SacI and SalI | SacI and SalI |
| pEMG-Δ*mqsRA* | Acc65I and SalI | Acc65I and SalI |
| pEMG-Δ*mazEF* | PvuII and BamHI | SacI* and BamHI |
| pEMG-Δ1716-1717 | BamHI and SacI | BamHI and SacI |
| pEMG-Δ*brnTA* | SacI and XbaI | SacI and XbaI |
| pEMG-Δ4151-4152 | EcoRI and KpnI | EcoRI and KpnI |
| pEMG-Δ*relBE* | BamHI and SacI | BamHI and SacI |
| pEMG-Δ*yefM-yoeB* | SacI and XbaI | SacI and XbaI |
| pEMG-Δ*relB*$_2$-*parE* | EcoRI and XbaI | EcoRI and XbaI |
| pEMG-Δ*xre* | Bpu1102I* and SacI | BamHI* and SacI |
| pEMG-Δ*higA* | SacI and Eco147I | SacI and SmaI |
| pEMG-Δ*hicB-1* | BamHI and EcoRI | BamHI and EcoRI |
| pEMG-Δ*higA*$_2$ | BstEII* and SacI | SmaI and SacI |
| pEMG-Δ*mqsA* | Mva1269I* and Acc65I | SmaI and Acc65I |
| pEMG-Δ*mazE* | PvuII and SalI | SalI and SmaI |
| pEMG-Δ1716 | EheI and SacI | SmaI and SacI |
| pEMG-Δ*brnA* | Psp1406I* and XbaI | SmaI and XbaI |
| pEMG-Δ4151 | EcoRI and PvuII | EcoRI and SmaI |
| pEMG-Δ*relB* | Eco91I* and BamHI | SmaI and BamHI |
| pEMG-Δ*yefM* | SacI and XbaI | SacI and XbaI |
| pEMG-Δ*relB*$_2$ | EcoRI and SacI | EcoRI and SacI |

[a] *DNA end cleaved with particular enzyme is blunt-ended with DNA Polymerase I Klenow Fragment
